# Comprehensive genome and transcriptome analysis reveals genetic basis for gene fusions in cancer

**DOI:** 10.1101/148684

**Authors:** Nuno A. Fonseca, Yao He, Liliana Greger, PCAWG3, Alvis Brazma, Zemin Zhang

**Affiliations:** European Molecular Biology Laboratory, European Bioinformatics Institute (EMBL-EBI), Wellcome Trust Genome Campus, Hinxton, Cambridge CB10 1SD, UK; Peking-Tsinghua Centre for Life Sciences, BIOPIC, and Beijing Advanced Innovation Centre for Genomics, Peking University, Beijing, 100871, China

## Abstract

Gene fusions are an important class of cancer-driving events with therapeutic and diagnostic values, yet their underlying genetic mechanisms have not been systematically characterized. Here by combining RNA and whole genome DNA sequencing data from 1188 donors across 27 cancer types we obtained a list of 3297 high-confidence tumour-specific gene fusions, 82% of which had structural variant (SV) support and 2372 of which were novel. Such a large collection of RNA and DNA alterations provides the first opportunity to systematically classify the gene fusions at a mechanistic level. While many could be explained by single SVs, numerous fusions involved series of structural rearrangements and thus are composite fusions. We discovered 75 fusions of a novel class of inter-chromosomal composite fusions, termed *bridged fusions*, in which a third genomic location bridged two different genes. In addition, we identified 522 fusions involving non-coding genes and 157 ORF-retaining fusions, in which the complete open reading frame of one gene was fused to the UTR region of another. Although only a small proportion (5%) of the discovered fusions were recurrent, we found a set of highly recurrent fusion partner genes, which exhibited strong 5’ or 3’ bias and were significantly enriched for cancer genes. Our findings broaden the view of the gene fusion landscape and reveal the general properties of genetic alterations underlying gene fusions for the first time.

---

The simultaneous availability of whole genome and RNA sequencing data for a large panel of International Cancer Genome Consortium (ICGC) tumour samples creates an unprecedented opportunity to identify novel gene fusions and understand their genetic basis. Gene fusions were detected by combining the output of two fusion discovery methods and genomic rearrangement (structural variant-SV) information, and several filters were implemented to exclude artefacts or those also present in the ICGC or GTEx normal samples [1] (Online Methods; Supplementary Fig 1; Supplementary Fig 2). Gene fusions were associated to the nearest SV with both breakpoints located within 250 kb of the two corresponding fusion breakpoints. These gene fusions were categorized based on novelty, recurrence, known oncogenic gene partners, breakpoint distance, breakpoint location, and matched SV support for downstream analyses (Fig. 1A).

**Figure 1.**
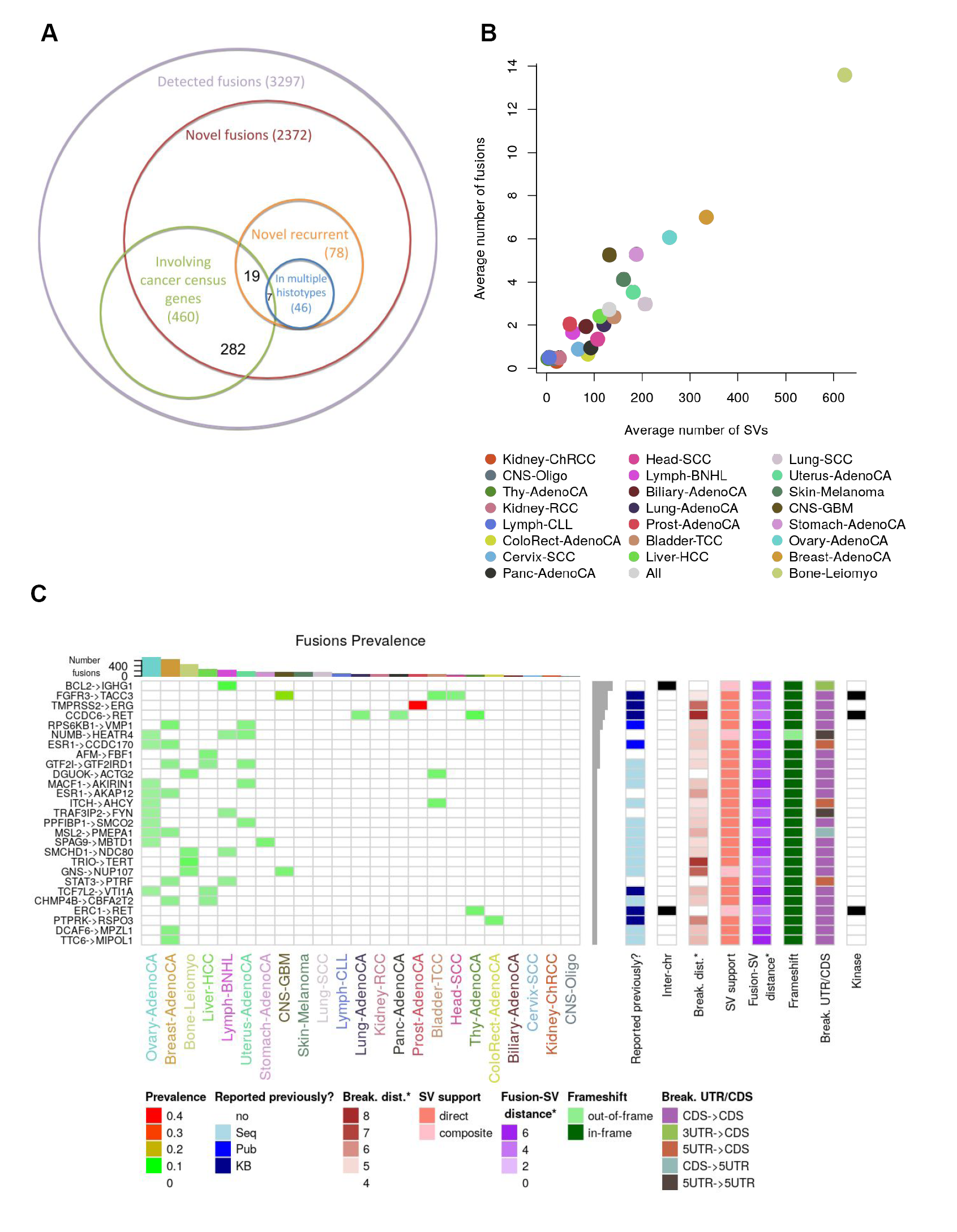
Gene fusion detected in 1185 donors. (A). The number of all detected and novel fusions and their overlap with the cancer census genes. Only a minority of the fusions are recurrent across samples, and over a half of the recurrent fusions are present in several cancer histotypes. From the novel recurrent fusions 19 involve cancer census genes and 7 of them are in multiple histotypes. (B). The number of fusions discovered per sample is highly uneven across cancer types, ranging from less than 1 to over 10 fusions per sample. The Pearson correlation between the number of fusions per sample and the number of SVs per sample is 0.96. (C). Features of the 27 most recurrent in-frame or ORF-retaining fusions. ChimerDB 3.0 [18] was used as a reference of previously reported gene fusions. The values with a star (‘*’) are log10. Kinase column indicates whether one of the gene partners is a kinase gene.

**Figure 2.**
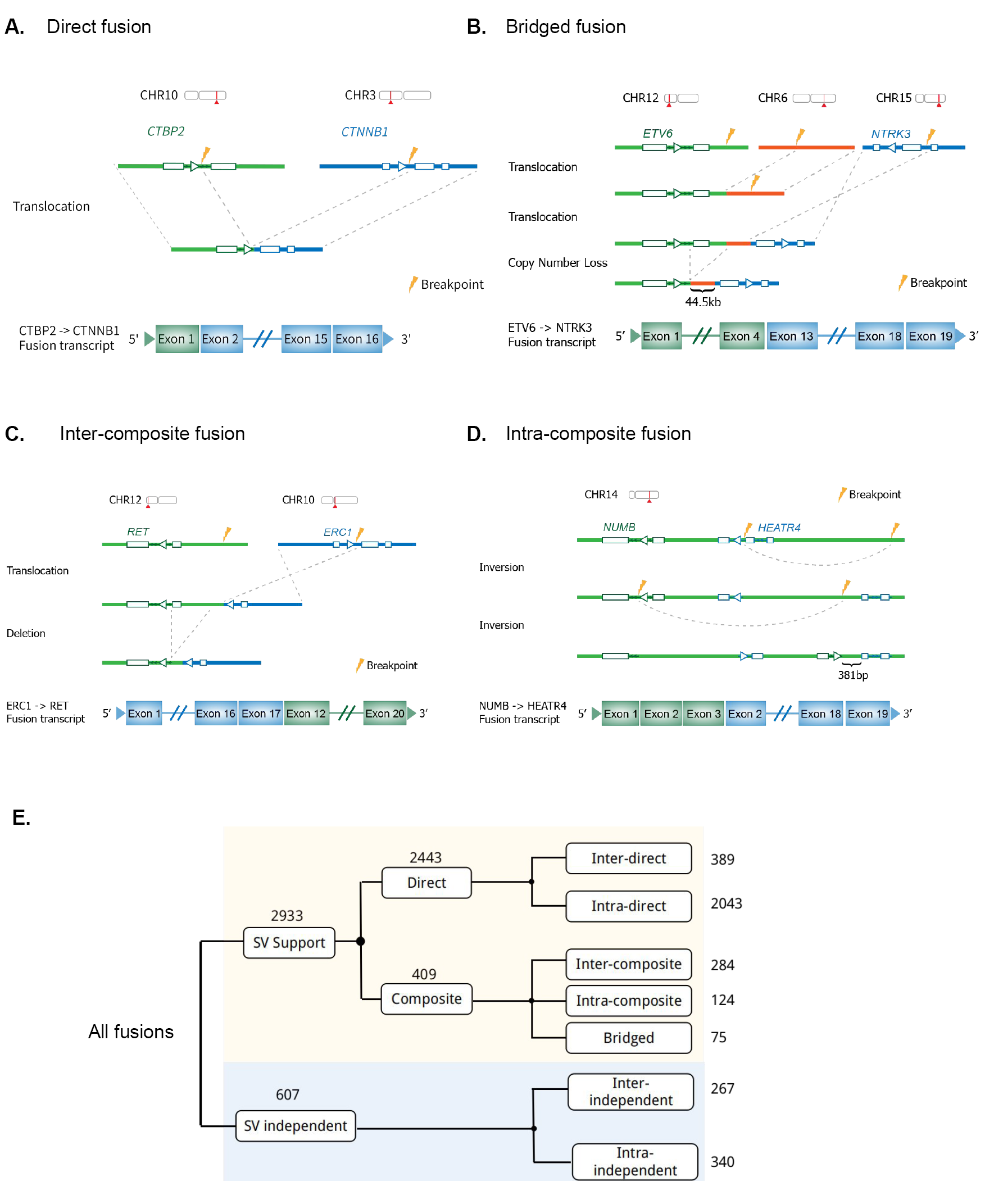
Fusion classification based on the underlying structural arrangements. A-D) Schematic representation of examples of different types of SV-supported fusions: A) direct fusions, B) bridged fusions, C) inter-composite fusions, and D) intra-composite fusions. Bridged fusions are those composite fusions formed by a third genomic segment bridging two different genes. Only one of the possible orders of genomic arrangement is depicted in each case, with breakpoints highlighted as thunderbolts. E) Systematic classification scheme of all gene fusions based on underlying SVs. Numbers of fusion events of different classes are shown to the right.

The average number of putative gene fusions per sample varies considerably across histological types (mean = 3, median = 2, sd = 3) and the number of gene fusions is highly correlated with the average number of SVs (Pearson correlation 0.96), supporting SVs as a major cause of gene fusions (Fig. 1B). For instance, bone-leimyocarcinomas harbour ∼14 fusions per sample on average, while 10 cancer types have less than one fusion per sample. The fusion frequencies were also associated with other types of alterations (Supplementary Fig. 3) but the observed correlation is lower (Supplementary Fig. 4).

**Figure 3.**
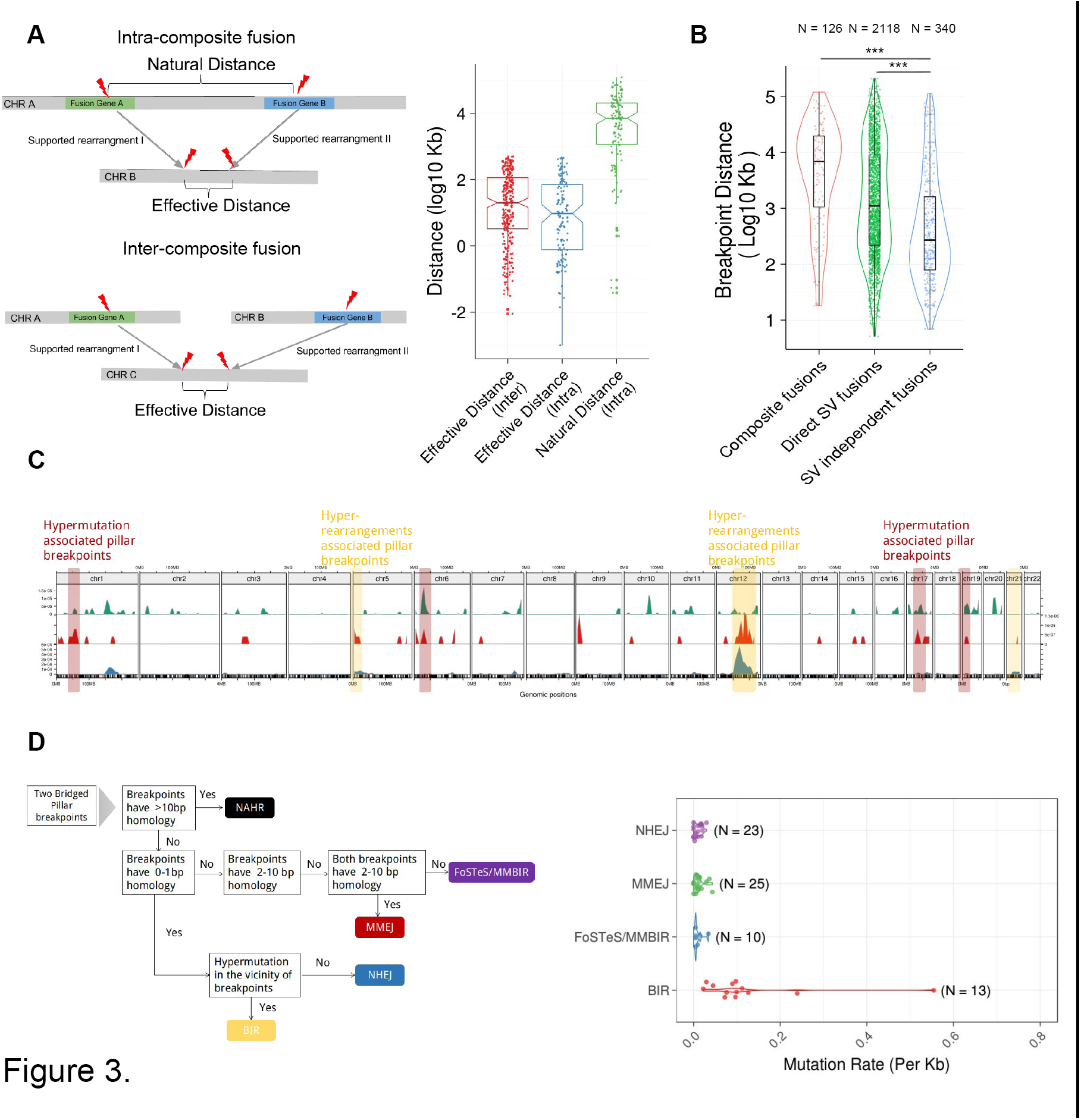
Genomic features of fusion-associated SVs. (A). Supported rearrangements for composite fusions bring the fused segments of two genes significantly closer. The cartoon on the left represents three different distances used in this comparison. Natural distance indicates the native distance between two related SV breakpoints. Effective distance indicates the distance between the final two breakpoints of the intra – composite / inter-composite fusions. B) The distribution of distances of fusion breakpoints among intra-chromosomal composite, direct SV-support and SV-independent fusions. P-value was determined by Student’s t test (***: P < 0.001). C) Bridged pillar breakpoints tend to be associated with genomic regions with hyper-rearrangements or hypermutations. The top track shows the mutation density above the genome-wide average mutation density for all aliquots containing bridged fusions, and the bottom track shows the density of all SV breakpoints above the genome-wide average SV density. The middle track shows the density of pillar breakpoints. D) The classification of bridged fusion formation mechanisms based on breakpoint features. The classification criteria are inferred primarily from [21], [22], [23], leading to the assignment of five types of mechanisms: non-allelic homologous recombination(NAHR), break-induced repair (BIR), non-homologous end joining (NHEJ), microhomology-mediated end joining (MMEJ) and template switching/microhomology-mediated break induced repair (FoSTeS / MMBIR). As expected, BIR-induced bridged regions show hypermutation status (based on the mutation rate in the vicinity of a 100kb window of bridged regions).

**Figure 4.**
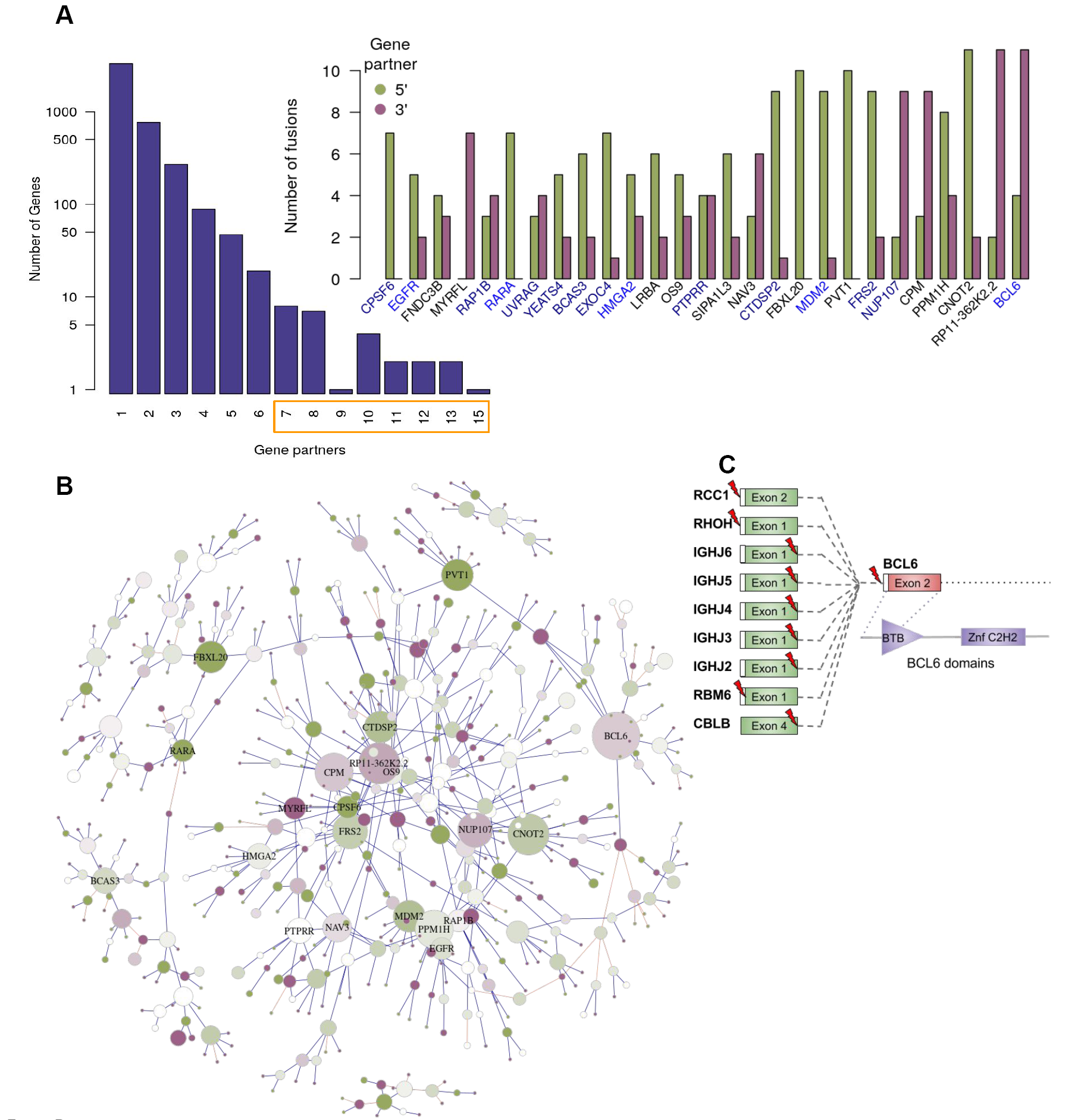
A) Left-although most genes appear as fusion partners with only with one gene, some of the genes may have up to 15 different fusion partners. Right-the most “promiscuous” fusion gene partners, with at least 7 partners and their preference as 5’ or 3’ partners. Genes in blue are known cancer genes (COSMIC v80) while genes in dark blue have reported interactions (in the STRING database [24]) with cancer genes. B) Connected clusters of at least 10 genes. Genes are represented as nodes and the size of a node is proportional to the number of gene fusion partners. Two nodes are connected if one fusion was detected involving the two genes: an edge is colored blue if the fusion has matched structural rearrangements evidence and is colored orange otherwise. Nodes and connections are only shown between promiscuous genes (non-promiscuous genes are not displayed). The color intensity indicates if a gene is involved more often in a fusion as 3’ (purple) or 5’ (green) gene or both (white). C) Schematic representation of the fusions involving *BCL6* as the 3’ partner and other protein coding genes as the 5’ partners. In all cases BCL6 protein domains remains intact.

By examining somatic rearrangement events and fusions simultaneously, we found 2618 fusion events that could be explained by single genomic rearrangements, with duplication as the predominant type, suggesting genomic amplification as a major source for gene fusions (Supplementary Fig. 5). Although 1288 of these rearrangements occurred within gene bodies by juxtaposing exons from two different genes, 720 other rearrangements disrupted neither of the fusion gene partners (Supplementary Fig. 5).

Notably, a large number of fusions, including known fusions (Supplementary Fig. 6) namely ETV6-NTRK3 [2] could not be associated with any single SV event. The ETV6-NTRK3 fusion was present in a head and neck thyroid carcinoma sample, linking exon 4 of ETV6 to exon 12 of NTRK3. We found three separate SVs: i) a translocation of ETV6 (chr12:12,099,706) to chromosome 6 (chr6:125,106,892); ii) a translocation of NTRK3 (chr15:88,694,049) also to chromosome 6 (chr6:125,062,387); and iii) an additional copy number loss (chr12:12,032,501-chr12:12,099,705) spanning from ETV6 intron 5 to the exact SV breakpoints (chr12:12,099,706), jointly bringing ETV6 within 45 kb upstream of NTRK3, a distance that would allow transcriptional read-through [3] or splicing [4] to yield the ETV6-NTRK3 fusion [5] (Fig. 2B). Thus, the short chromosome 6 segment appeared to function as a bridge, linking two other genomic locations to facilitate a gene fusion. We term such products bridged fusions. This novel class of fusions are not uncommon. Out of a total of 436 fusions supported by two separate SVs, 75 are bridged fusions. The lengths of bridge regions ranges from 1bp to 496 kb, with a median size of 3.7 kb and an average size of 62.96 kb (Supplementary Fig. 7). One third of the bridge regions are located in genic regions and none of such genic regions were found to be involved in other fusions in the corresponding samples.

Aside from bridged fusions, 344 additional fusions are supported by more than one SV. These multi-SV fusions are collectively termed *composite fusions*. For example, the known *ERC1-RET* fusion, previously described as a product of the translocation t(10;12)(q11;p13), was detected in two samples of head and neck and thyroid carcinomas. While there was no evidence of direct translocation in either case, they were both supported by an inter-chromosomal translocation and an intra-chromosomal rearrangement (deletion or inversion), resulting in the connection of either exon 12 or exon 17 of *ERC1* to the exon 12 of *RET* (Fig. 2C).

While fusion transcripts formed by two adjacent genes are often viewed as non-genomic transcription-induced chimeras [3], we observed that such chimera formation could possibly be facilitated by composite DNA rearrangements. For one of the tumours with the recurrent *NUMB-HEATR4* fusion, we detected two consecutive inversions, bringing the *NUMB* exon 3 within 381 bp of the *HEATR4* exon 2 (Fig. 2D). The much shorter distance, down from the natural distance of 14 kb, would allow fusion formation by splicing. Such fusion could also result from duplication – it was supported by a duplication spanning from exon 2 of the *NUMB* gene to exon 4 of the *HEATR4* gene in two other tumours.

Based on the nature of underlying genomic rearrangements, we propose a unified fusion classification system (Fig. 2E). Overall, we identified 75 bridged fusions, 284 inter-composite fusions generated by a translocation linking two genes from different chromosomes followed by a second intra-chromosomal rearrangement, and 125 intra-composite fusions generated by multiple intra-chromosomal rearrangements.

To further understand the mechanisms of composite fusion formation, we examined various features associated with such fusions. Intra-composite fusion partners were brought significantly closer to each other, from the median natural distance of 6,836 kb to the median of 7.9 kb (Wilcoxon Rank Sum Test, P < 2.2e-16, Fig. 3A). Inter-composite fusion partners also exhibited similarly short gene distances post-translocation (Fig. 3A). Although bridged fusion pillar breakpoints were distributed broadly across the genome with no particular hotspots (Supplementary Fig. 8), they were over-represented on chromosome 12 (Fig. 3C, Supplementary Fig. 9). Notably, such pillar breakpoints were associated with genomic regions with high mutation density or high rearrangement density (Fig. 3C), indicating the connection with mutational processes such as chromothripsis and kataegis [6,7] (Fig. 3C). Eighty eight of the 150 pillar breakpoints overlapped with chromothripsis regions [25], and 33 of these 88 regions further accompanied kataegis [8]. For the remaining 62 pillar breakpoints, while we did not find local enrichment for any specific sequence motifs or fragile sites, 53% of these were located within the *Alu* and *LINE* repeat elements or other simple repeats. Based on the presence of microhomology in the breakpoints and the hypermutation status of bridge regions, we inferred that 23 of the 75 bridged fusions were formed by replication-based DNA double strand repair process, and 52 fusions were formed by either non-homologous or microhomology-mediated end-joining (*NHEJ/MMEJ*) (Fig. 3D). Thus, multiple mechanisms could lead to bridged fusions, with chromothripsis playing a key role.

While most fusions had direct or composite SV support, the remaining 18%, including known fusions like *RHOH-BCL6* [9] did not have obvious SV evidence. Thus, either these genes were fused directly at the RNA level or the underlying supporting SVs were somehow missed. The latter was evidenced by an observation that known fusions, such as *TMPRSS2-ERG* [10], did not have consistent SV support in all samples where it was detected (in 4 out of 6 samples *TMPRSS2-ERG* fusion was supported by a deletion, while in the other two samples it did not have any SV support). On the other hand, RNA read-through events were also likely: the 340 SV independent, intra-chromosomal fusions had significantly closer breakpoints than those with direct or composite SV support (Fig. 3B). Read-throughs for such close-by genes have been observed previously [5].

We next examined the nature of the fusion products. While ∼36% of all detected fusion transcripts were predicted to be in-frame, there were other types of fusions involving UTRs, non-coding genes and other combinations (Supplementary Fig. 10). In particular, certain UTR-mediated fusion transcripts preserve complete coding sequences of one fusion partner. These include a known fusion *TBL1XR1–PIK3CA* in a breast tumour, in which the 3’ complete *PIK3CA* ORF is overexpressed by the promoter of the 5’ *TBL1XR1* [11]. We found 6 additional novel ORF-retaining fusions, specifically *RCC1-BCL6, RP11-362K2.2-CDK4, CTBP2-CTNNB1*, *RBM6-BCL6*, *SF3B3-CDH1* and *RNF38-PDCD1LG2*, in which the complete ORF of cancer-related genes were the 3’ partners, benefiting from the stronger promoters of the 5’ partners. Notably, the oncogene *CTNNB1*, not previously known to be activated through gene fusion, showed elevated expression in the fusion-containing gastric tumour, at level similar to those with *CTNNB1* amplification, indicating that gene fusions could drive *CTNNB1* overexpression (Supplementary Fig. 11).

Of 44 recurrent in-frame or ORF-retaining fusions (Fig. 1C), 31 have been previously well described, including *CCDC6-RET, ERC1-RET, FGFR3-TACC3, TRIO-TERT, RPS6KB1-VMP1, SEC61G-EGFR*, and *TMPRSS2-ERG*. A less characterized recurrent fusion, *DCAF6-MPZL1*, is ORF-retaining and recurrent in breast tumours. *MPZL1* encodes a cell adhesion surface receptor, with its overexpression linked to cancer cell migration in hepatocellular carcinoma and response to trastuzumab in breast cancer [12,13]. Based on combined expression and DNA copy number analysis, the high expression of MPZL1 in fusion-positive samples was likely driven by gene fusions instead of DNA copy number gain (Supplementary Fig.12). Also of interest is the *GNS-NUP107* intra-composite, in-frame fusions found in sarcoma and glioblastoma samples, with the exon 9 of GNS fused to either exon 20 or exon 9 of *NUP107* by an inversion and a deletion, retaining the *NUP107* leucine zipper motif in its carboxyl-terminal region (Supplementary Fig. 13). *NUP107* is known to be overexpressed in breast cancer, with association with poor prognosis [14] and anti-apoptotic functions in astrocytoma [15]. The *NUP107* expression in the fusion-containing samples was elevated, consistent with *GNS-NUP107* as a possible oncogenic driver event.

Only 3% of the 2373 novel fusions were recurrent (Fig. 1A), with the majority occurring only in one histotype, while 7 were found across multiple histotypes. Of the 12 most recurrent gene fusions (Supplementary Fig. 14), 2 have been previously described (*CCDC6-RET* [16], *FGFR3-TACC3* [17] and 6 detected in *TCGA* [18]), while 4 were novel. By contrast, many of the fusion partner genes were highly recurrent. Although 3294 of the 4515 unique genes involved in fusions had only one fusion partner, 35 genes had more than five partners (Fig. 4A). Most of these “promiscuous” genes tended to be selective in being either a 5’ or 3’ partner (Fig. 4A). Moreover the set was overrepresented in cancer census genes (one tailed Fisher’s exact test, odds ratio (OR) = 8.66, P < 1.1e-15) and in PCAWG’s cancer driver genes (one tailed Fisher’s exact test, odds ratio (OR) = 12.27, P < 2.2e-16).

To further investigate the promiscuous fusion gene partners, we constructed a network by connecting any two genes that were detected as fused in at least one sample (Supplementary Fig. 15). Most genes belonged to small clusters but several larger clusters emerged (Supplementary Fig. 16). Focusing on clusters with at least 10 genes (Fig. 4B), we found that they were significantly enriched in several cancer-related pathways (Supplementary Fig. 17, Benjamini-Hochberg corrected p-value cut-off of 0.01) and in protein-protein interactions (Supplementary Fig. 18). For example, *BCL6*, a known oncogene, was involved in 15 different fusions, mostly as a 3’ partner, and had the breakpoints conserved (*RHOH*, *IGHJ6/5/4/3/2*, *RCC1*, *RP11-731F5.1*, *RBM6* and *CBLB*). All these *BCL6* fusions contained the intact exon 2 of *BCL6* and seemed to co-opt the regulatory sequences of the 5’ fusion partners that replace the 5’ untranslated region of the *BCL6* (Fig. 4C). This pattern had been reported previously in primary gastric high-grade B-cell lymphoma [19,20]. In general, the breakpoints and their positions (3’ or 5’) were often conserved in promiscuous genes and did not show association with other genomic feature such as common fragile sites (Supplementary Fig. 20), indicating that these genes tend to selectively fuse to other genes. Taken together the data suggests that at least some of the promiscuous fusion partners might play a functional role in cancer progression.

In conclusion, this is the first study that systematically compares and integrates gene fusions with whole genome rearrangements across many tumour types. We discovered thousands of high-confidence cancer-specific fusions, developed a systematic classification of fusion events, and proposed a novel bridged fusion mechanism to explain how genome rearrangements can lead to a gene fusion. It is worth noting that although only a minority (149, 5%) of the discovered fusions were recurrent, thus indicating that many fusions are likely to be “passengers”, the promiscuous fusion gene partners were often linked to cancer related pathways, thereby indicating a possible functional role in cancer. Our findings broaden the view of the gene fusion landscape in human cancers and reveal the general properties of genetic alterations underlying gene fusions for the first time.

